# The impact of RSV/SARS-CoV-2 co-infection on clinical disease and viral replication: insights from a BALB/c mouse model

**DOI:** 10.1101/2023.05.24.542043

**Authors:** Dorothea R. Morris, Yue Qu, Kerrie S. Thomason, Aline Haas de Mello, Richard Preble, Vineet D. Menachery, Antonella Casola, Roberto P. Garofalo

## Abstract

RSV and SARS-CoV-2 are prone to co-infection with other respiratory viruses. In this study, we use RSV/SARS-CoV-2 co-infection to evaluate changes to clinical disease and viral replication in vivo. To consider the severity of RSV infection, effect of sequential infection, and the impact of infection timing, mice were co-infected with varying doses and timing. Compared with a single infection of RSV or SARS-CoV-2, the co-infection of RSV/SARS-CoV-2 and the primary infection of RSV followed by SARS-CoV-2 results in protection from SARS-CoV-2-induced clinical disease and reduces SARS-CoV-2 replication. Co-infection also augmented RSV replication at early timepoints with only the low dose. Additionally, the sequential infection of RSV followed by SARS-CoV-2 led to improved RSV clearance regardless of viral load. However, SARS-CoV-2 infection followed by RSV results in enhanced SARS-CoV-2-induced disease while protecting from RSV-induced disease. SARS-CoV-2/RSV sequential infection also reduced RSV replication in the lung tissue, regardless of viral load. Collectively, these data suggest that RSV and SARS-CoV-2 co-infection may afford protection from or enhancement of disease based on variation in infection timing, viral infection order, and/or viral dose. In the pediatric population, understanding these infection dynamics will be critical to treat patients and mitigate disease outcomes.

**Author Summary:** Infants and young children are commonly affected by respiratory viral co-infections. While RSV and SARS-CoV-2 are two of the most prevalent respiratory viruses, their co-infection rate in children remains surprisingly low. In this study, we investigate the impact of RSV/SARS-CoV-2 co-infection on clinical disease and viral replication using an animal model. The findings indicate that RSV infection either simultaneously or prior to SARS-CoV-2 infection in mice protect against SARS-CoV-2-induced clinical disease and viral replication. On the other hand, infection with SARS-CoV-2 followed by RSV results in worsening of SARS-CoV-2-induced clinical disease, but also protection from RSV-induced clinical disease. These results highlight a protective role for RSV exposure, given this occurs before infection with SARS-CoV-2. This knowledge could help guide vaccine recommendations in children and sets a basis for future mechanistic studies.

**Graphical Abstract:** 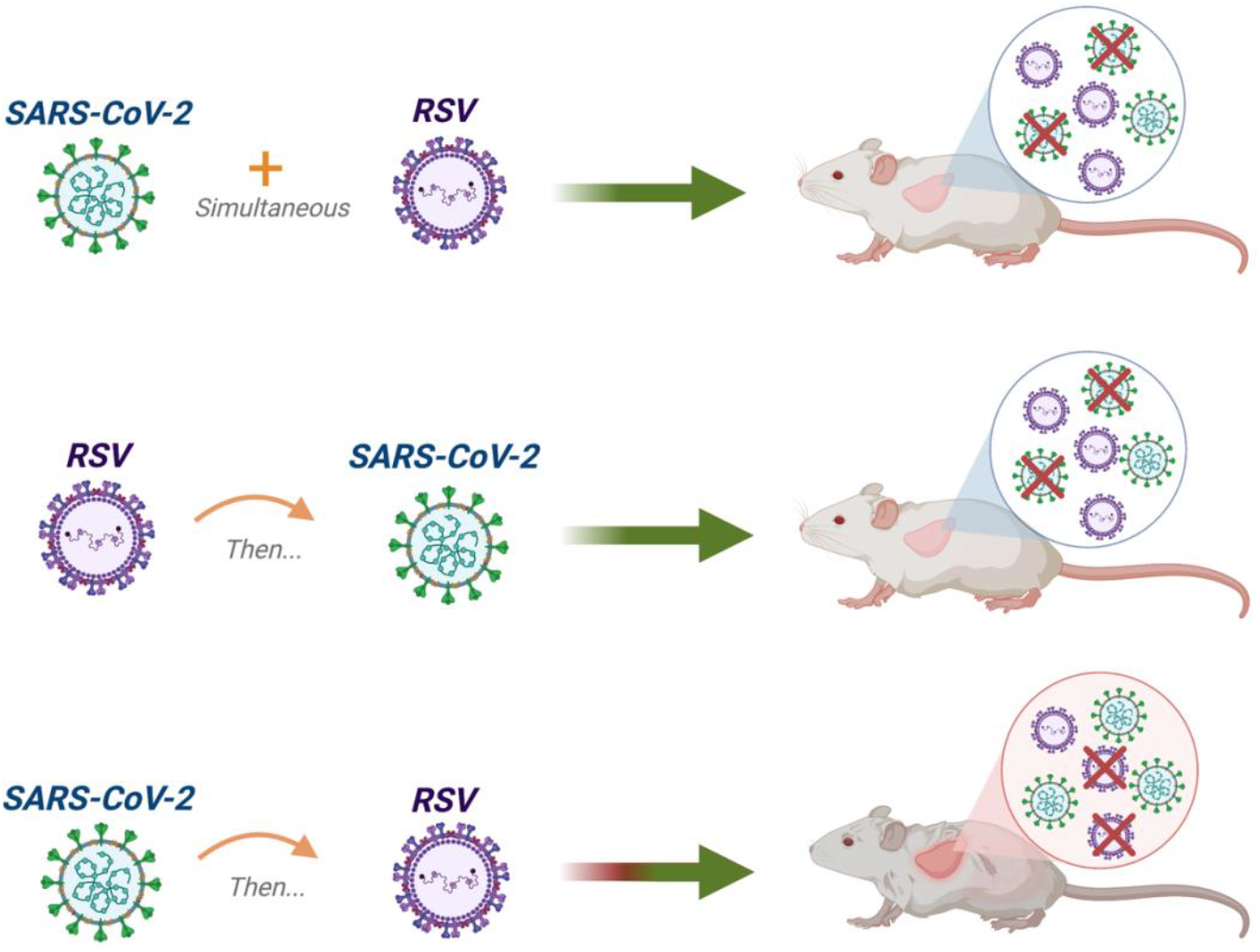

## Introduction

Respiratory syncytial virus (RSV) is the leading cause of respiratory illness in infants and young children with an estimated 33 million infections each year [1]. During the COVID-19 pandemic, cases of severe acute respiratory syndrome coronavirus-2 (SARS-CoV-2) in children <5 years of age had remained low until the emergence of the Delta (B.1.617.2) and Omicron (B.1.1.529) variants [2]. According to the American Academy of Pediatrics, there have been a total of 15.5 million SARS-CoV-2 infections in children since the onset of the pandemic and ∼65% of these cases were recorded between September 2021 and March 2023. Additionally, the presence of SARS-CoV-2 has placed unique epidemiological pressures on all common respiratory viruses, leading to a persistent circulation of RSV in the population since May 2021 [3, 4]. This has increased the likelihood of co-infections between RSV and SARS-CoV-2 among children. Interestingly, children have been found to be particularly susceptible to respiratory viral co-infection, and be more likely to be co-infected with SARS-CoV-2 than adults [5-7].

Despite this, review of clinical findings for RSV/SARS-CoV-2 co-infection have reported rates on average of 3% [8-10]. These patients require moderately more supplemental care when hospitalized, but none were found to have an increased risk of mortality when compared to either virus separately. These findings are surprising as the transmission rate of RSV or SARS-CoV-2 in children is high [7, 11, 12]. Additionally, the inflammatory milieu of RSV or SARS-CoV-2 would suggest co-infection between these two viruses could result in significantly worse disease outcomes, like that of influenza/SARS-CoV-2 co-infection [13-15].

The phenomenon of viral-viral co-infection is still poorly understood. Given the high rate of transmission and the potential severity of disease, the objective of this study was to characterize baseline alteration in clinical disease and viral replication following RSV/SARS-CoV-2 co-infection using a BALB/c mouse model. We take into consideration the severity of RSV infection, the sequential effect of infection, and the impact of timing on infection. To do so, mice were inoculated with a low or high dose of RSV, were infected with RSV followed by SARS-CoV-2 or SARS-CoV-2 followed by RSV, and were either infected simultaneously or 48 hours after primary inoculation. In summary, we find that the simultaneous co-infection of RSV/SARS-CoV-2 and the infection of RSV followed by SARS-CoV-2 results in protection from SARS-CoV-2-induced clinical disease and viral replication. Reciprocally, mice infected with SARS-CoV-2 followed by RSV have protection from RSV-induced disease and viral replication in the lung, but exhibit worsening of SARS-CoV-2-induced disease. Collectively, these data suggest that the timing, order, and dose of virus during co-infection can exacerbate or diminish disease. Given this, our findings shed light on an important public health concern and provide baseline data to inform treatment and future mechanistic studies.

## Results

### Co-infection of RSV and SARS-CoV-2 protects against SARS-CoV-2 induced disease

To investigate whether simultaneous co-infection of SARS-CoV-2 with RSV affects clinical disease, groups of BALB/c mice (n=10) were intranasally inoculated with 1×10^6^ 50% tissue culture infective dose (TCID_50_/mL) of CMA3p20 (mouse-adapted SARS-CoV-2) mixed with 2.5×10^6^ plaque forming units (PFU) or 1×10^7^ PFU of RSV Long Strain in 50uL of PBS (Figure 1A). We found that the infection of SARS-CoV-2 combined with the lower infectious dose of RSV results in protection from SARS-CoV-2 induced bodyweight loss with a maximum improvement of 7.9% at day 3 p.i. (Figure 1B). These co-infected mice also had significant improvements in illness as compared to the SARS-CoV-2/PBS control mice (Figure 1C). Mice infected with SARS-CoV-2 combined with the higher infectious dose of RSV resulted a bodyweight loss and illness score pattern identical to that of the RSV/PBS control mice, indicating no further worsening of clinical disease due to the presence of SARS-CoV-2 (Figure 1D and 1E). These results suggest that the simultaneous infection of RSV with SARS-CoV-2 protects against SARS-CoV-2 induced disease, regardless of the RSV infection load.

**Fig. 1.**
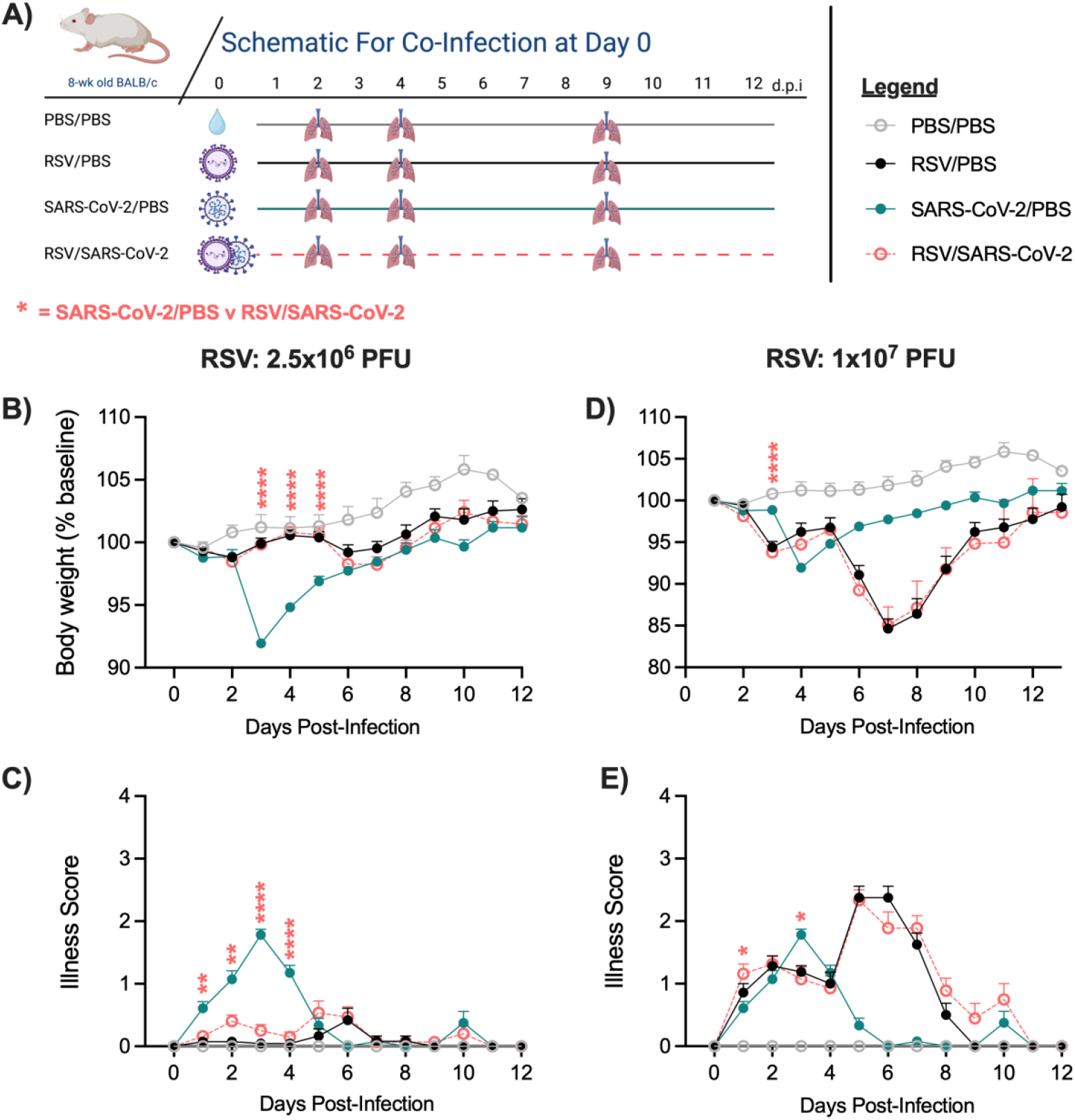
Assessment of clinical disease in BALB/c mice simultaneously co-infected with RSV and SARS-CoV-2. (A) The experimental design for Figure 1 and Figure 2. BALB/c mice were IN inoculated with PBS, RSV/PBS at a dose of either 2.5×10^6^ or 1×10^7^ PFU, SARS-CoV-2/PBS at a dose of 1×10^6^ TCID_50_/mL, or RSV/SARS-CoV-2 simultaneously. (B-E) Mice were monitored for changes in bodyweight loss and illness score over the 12-day infection period. Data are pooled from three independent experiments for mice infected with the RSV dose of 2.5×10^6^ PFU (n ≤ 25 mice/group). Data are pooled from two independent experiments for mice infected with the RSV dose of 1×10^7^ PFU (n=20 mice/group). Data are expressed as mean ± SEM. Significant results as compared to the SARS-CoV-2/PBS control are marked with asterisks (* p ≤ 0.05, ** p ≤ 0.01, **** p ≤ 0.0001).

### Simultaneous co-infection of RSV and SARS-CoV-2 reduces SARS-CoV-2 replication

We next assessed if the simultaneous co-infection of RSV and SARS-CoV-2 would alter viral replication. Lung tissue was collected for RT-qPCR at the indicated time points corresponding to peak viral replication for either virus or a later time point to assess viral clearance (Figure 1A). Mice that were co-infected with the lower infectious dose of RSV demonstrated a significant fold increase of 4.25 in the RSV N gene copy number at day 2 p.i. as compared to the RSV/PBS control mice (Figure 2A). This increase in RSV N gene copy number was not appreciated at days 4 or 9 p.i. (Figure 2B, 2C). Interestingly, these same co-infected mice demonstrated a near complete reduction in SARS-CoV-2 N gene copy number at days 2, 4, and 9 p.i. (Figure 2A-2C). Mice that were co-infected with the higher infectious dose of RSV demonstrated no significant changes to RSV N gene copy number as compared to the RSV/PBS control mice at any time point assessed. Alternatively, these same co-infected mice demonstrated significant reductions in SARS-CoV-2 N gene copy number at days 2 and 4 p.i. (Figure 2D-2F). No co-infected mice showed signs of altered viral clearance from the lung tissue as compared to the appropriate controls (Figure 2C, 2F). These data suggest that RSV infection protects against SARS-CoV-2 replication during simultaneous co-infection, even at low infectious doses of RSV.

**Fig. 2:**
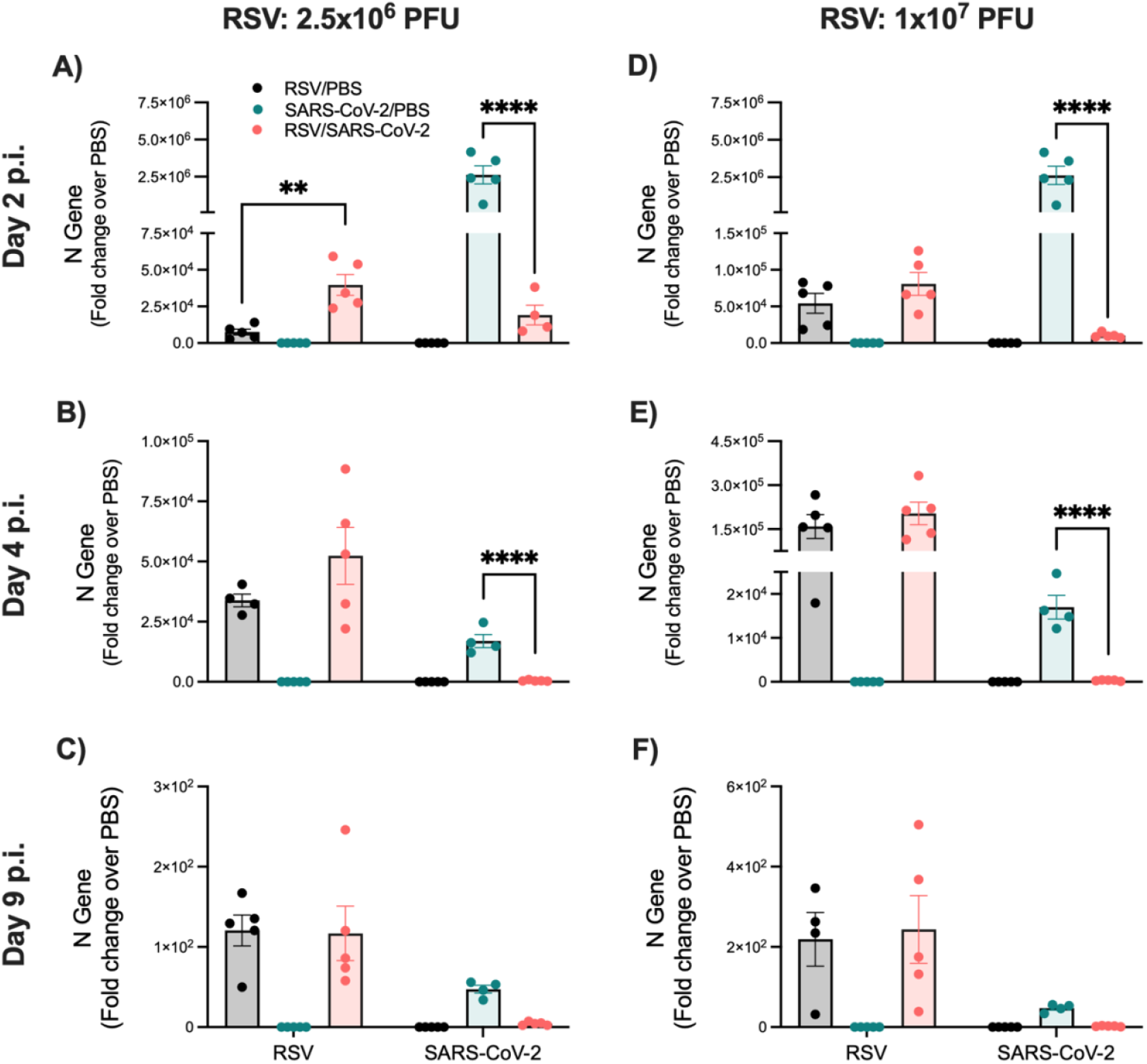
Assessment of gene expression by RT-qPCR in the lung of BALB/c mice simultaneously co-infected with RSV and SARS-CoV-2. Mice were infected as shown in Figure 1A. Lung tissue was collected at (A,D) day 2, (B,E) day 4, and (C,F) day 9 p.i. to assess RSV N and SARS-CoV-2 N gene expression by RT-qPCR. Data are expressed as mean ± SEM. Significant results as compared to the respective controls are marked with asterisks (** p ≤ 0.01, **** p ≤ 0.0001).

### Infection with SARS-CoV-2 followed by RSV results in worsening clinical disease

Though the simultaneous infection with two respiratory viruses is probable, it is more likely that a person would become co-infected over a short period of time. To simulate this, we designed a model in which BALB/c mice were intranasally inoculated with the secondary virus 48 h after the primary infection (Figure 3A). The groups include a mock control (PBS/PBS), RSV/PBS (2.5×10^6^ or 1×10^7^ PFU), SARS-CoV-2/PBS (1×10^6^ TCID_50_/mL), RSV infection followed by SARS-CoV-2 infection (RSV/SARS-CoV-2), and SARS-CoV-2 infection followed by RSV infection (SARS-CoV-2/RSV). We first assessed impacts to clinical disease in mice infected with RSV followed by SARS-CoV-2, including bodyweight loss and illness score. RSV/SARS-CoV-2 mice that received the lower infectious dose of RSV demonstrated protection from SARS-CoV-2 induced bodyweight loss with a maximum improvement of 7.41% at day 3 p.i. (Figure 3B). These mice also had significant improvements in illness as compared to the SARS-CoV-2/PBS control mice (Figure 3C). For RSV/SARS-CoV-2 mice that received the higher infectious dose of RSV, the bodyweight loss followed the same pattern as the RSV/PBS control mice (Figure 3D). These RSV/SARS-CoV-2 mice displayed a general trend towards worsening illness with significantly worse illness at days 7 and 8 p.i. as compared to the RSV/PBS control mice (Figure 3E). These data demonstrate that primary infection with RSV followed by SARS-CoV-2 infection protects from SARS-CoV-2 induced clinical disease, even at the lower infectious dose of RSV.

**Fig. 3.**
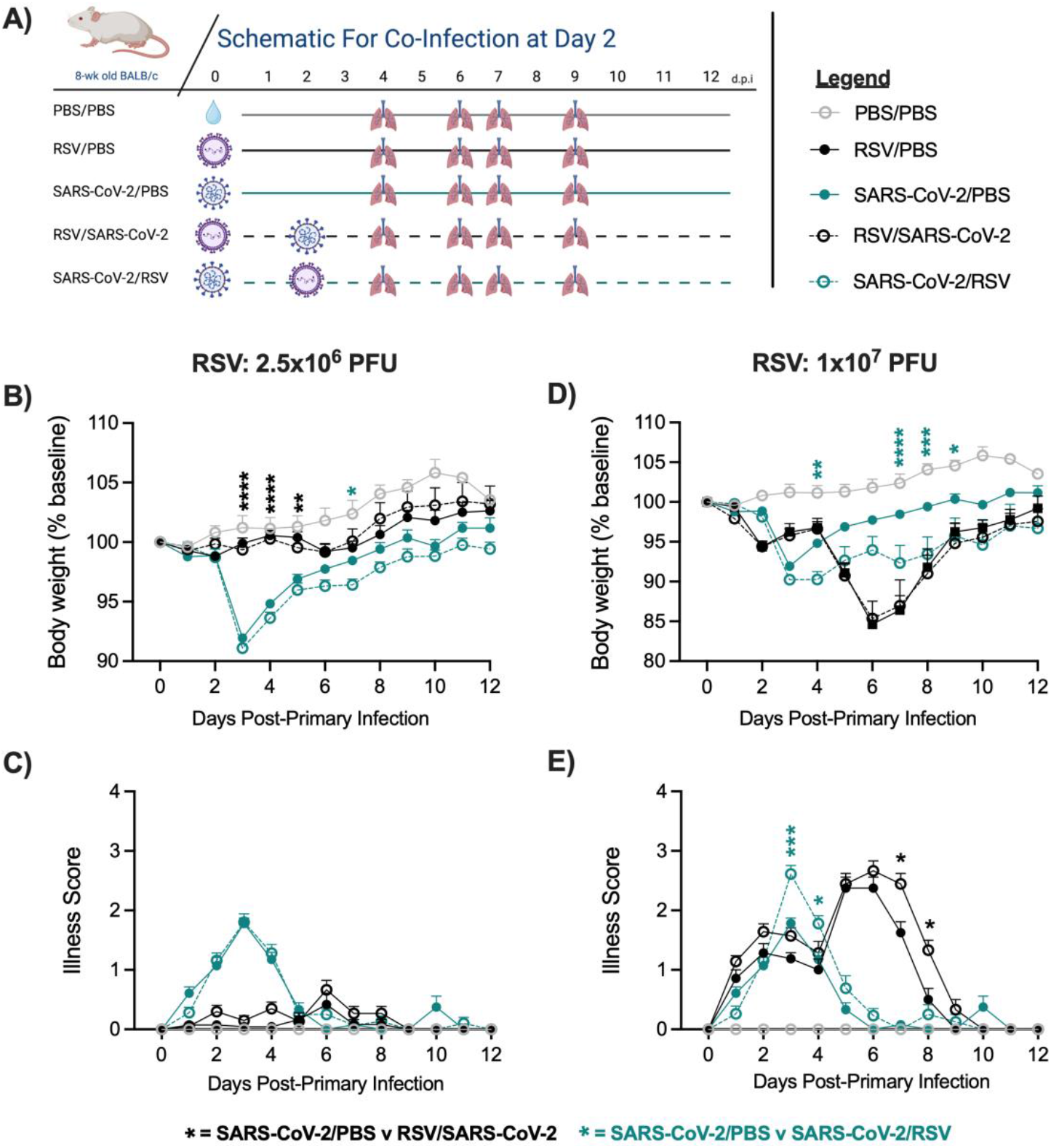
Assessment of clinical disease in BALB/c mice co-infected 48 h after primary infection. (A) The experimental design for Figures 3-5. BALB/c mice were IN inoculated with PBS, RSV/PBS at a dose of either 2.5×10^6^ or 1×10^7^ PFU, SARS-CoV-2/PBS at a dose of 1×10^6^ TCID_50_/mL, RSV followed by SARS-CoV-2, or SARS-CoV-2 followed by RSV. (B-E) Mice were monitored for changes in bodyweight loss and illness score over the 12-day infection period. Data are pooled from three independent experiments for mice infected with the RSV dose of 2.5×10^6^ PFU (n ≤ 25 mice/group). Data are pooled from two independent experiments for mice infected with the RSV dose of 1×10^7^ PFU (n ≤ 15 mice/group). Data are expressed as mean ± SEM. Significant results as compared to the RSV/PBS control are marked with black asterisks. Significant results as compared to the SARS-CoV-2/PBS control are marked with green asterisks (* p ≤ 0.05, ** p ≤ 0.01, *** p ≤ 0.001, **** p ≤ 0.0001).

We next assessed clinical disease in mice infected with SARS-CoV-2 followed by RSV. SARS-CoV-2/RSV mice that received the lower infectious dose of RSV demonstrated a trend towards worsening bodyweight loss, though this was only significant at day 7 p.i. (Figure 3B). There was no significant difference in illness score as compared to the SARS-CoV-2/PBS control mice (Figure 3C). For SARS-CoV-2/RSV mice that received the higher infectious dose of RSV, the bodyweight loss was significantly worse than the SARS-CoV-2/PBS control mice beginning at day 4 p.i. and continuing through day 12 p.i. (Figure 3D). These co-infected mice were also noted to have significantly worse illness as compared to the SARS-CoV-2/PBS control mice (Figure 3E).

Interestingly, these co-infected mice failed to display the strong double-weight loss curve elicited by the RSV/PBS control mice (Figure 3D). These data would indicate that primary infection with SARS-CoV-2 infection followed by RSV infection results in an exaggeration of SARS-CoV-2-induced clinical disease while protecting from RSV-induced disease.

### Primary infection with either RSV or SARS-CoV-2 reduces the replication of the corresponding secondary virus

To study the effects on viral replication, lung tissue was collected at timepoints corresponding to peak viral replication or viral clearances of the secondary infection, and compared to time-matched single infection controls (Figure 3A). We first examined the RSV/SARS-CoV-2 groups. Mice infected with the lower infectious dose of RSV followed by SARS-CoV-2 had no significant alteration to RSV N gene copy number at day 4 post-RSV infection as compared to the RSV/PBS mice (Figure 4A). Interestingly, the RSV N copy number was significantly reduced in the RSV/SARS-CoV-2 mice as compared to the RSV/PBS control mice at day 9-post RSV infection, indicating more efficient clearance of RSV from the lung tissue (Figure 4B). These same co-infected mice exhibited significantly lower SARS-CoV-2 N gene copy numbers at day 2 post-SARS-CoV-2 infection compared to the SARS-CoV-2/PBS control mice (Figure 4A). Similarly, mice infected with the higher infectious dose of RSV followed by SARS-CoV-2 had no alteration to RSV N gene copy numbers at day 4 p.i., but exhibited significantly reduced copy numbers at day 9 p.i. as compared to the RSV/PBS mice (Figure 4C, 4D). At peak viral replication for SARS-CoV-2, these same mice had significantly reduced SARS-CoV-2 N gene copy numbers as compared to the SARS-CoV-2/PBS control mice (Figure 4C). No significant changes to viral clearance were noted for either virus (Figure 4B, 4D). These findings indicate that primary infection with RSV followed by SARS-CoV-2 leads to reduced SARS-CoV-2 replication and increased RSV clearance in the lungs of BALB/c mice, regardless of the RSV infectious dose. This is similar to the simultaneous co-infection model in that SARS-CoV-2 replication is reduced, but the improved clearance of RSV appears to be unique to the to the sequential infection of RSV followed by SARS-CoV-2.

**Fig. 4:**
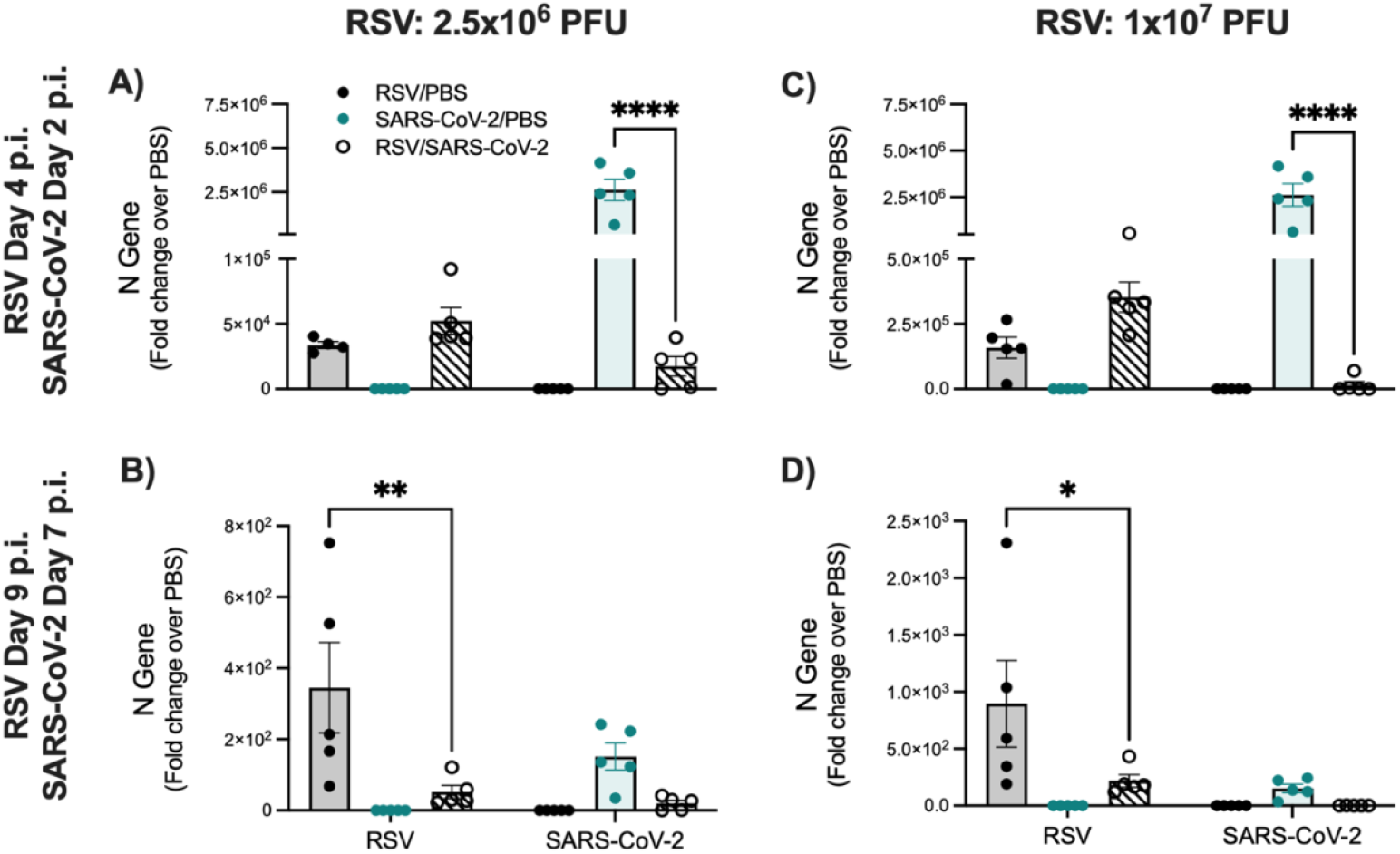
Assessment of gene expression by RT-qPCR in the lung of BALB/c mice infected with RSV followed by SARS-CoV-2. Mice were infected as shown in Figure 3A. Lung tissue was collected at (A,C) day 4 post-RSV infection/day 2 post-SARS-CoV-2 infection, and (B,D) day 9 post-RSV infection/day 7 post-SARS-CoV-2 infection to assess RSV N and SARS-CoV-2 N gene expression by RT-qPCR. Data are expressed as mean ± SEM. Significant results as compared to the respective controls are marked with asterisks (* p ≤ 0.05, ** p ≤ 0.01, **** p ≤ 0.0001).

Next, we assessed viral replication in the lung tissue of the SARS-CoV-2/RSV groups. Mice infected with SARS-CoV-2 followed by the lower infectious dose of RSV had significantly reduced RSV N gene copy numbers at days 2 and 4 post-RSV infection as compared to RSV/PBS mice (Figure 5A, 5B). In these same mice, no significant alteration to SARS-CoV-2 N gene copy number was appreciated at days 4, 6, or 9 post-SARS-CoV-2 infection as compared to the SARS-CoV-2/PBS mice (Figure 5A, 5B, 5C). Similarly, mice that received SARS-CoV-2 followed by the higher infectious dose of RSV, had significantly reduced RSV N gene copy numbers at days 2 and 4 post-RSV infection as compared to RSV/PBS mice (Figure 5D, 5E). In these same co-infected mice, no significant alteration to SARS-CoV-2 N gene copy number was appreciated at days 4, 6, or 9 post-SARS-CoV-2 infection as compared to the SARS-CoV-2/PBS mice (Figure 5D, 5E, 5F). These data suggest that primary infection with SARS-CoV-2 followed by RSV effectively reduces RSV replication in the lung of BALB/c mice, regardless of RSV infectious dose. This differs from the simultaneous co-infection model in that RSV replication was not altered. This change in RSV replication appears to be unique to the sequential infection of SARS-CoV-2 followed by RSV.

## Discussion

Despite high rates of transmission for RSV and SARS-CoV-2, the detection of RSV/SARS-COV-2 co-infections has remained low [16]. To better understand the co-infection dynamics of these two viruses, we have established baseline characteristics of clinical disease and viral replication using a BALB/c mouse model of RSV and SARS-CoV-2 co-infection. We take into consideration the severity of RSV infection, the sequential effect of infection, and the impact of timing on infection. To our knowledge, this is the first detailed description of an RSV/SARS-CoV-2 co-infection in an animal model. Here, we find that the simultaneous exposure of RSV and SARS-CoV-2 results in protection from SARS-CoV-2 induced clinical disease and viral replication, even in mice that received the lower infectious dose of RSV. Mice that received the primary infection of RSV followed by SARS-CoV-2 had similar outcomes to that of the simultaneous exposure, demonstrating protection from SARS-CoV-2 induced disease and reductions in SARS-CoV-2 replication. In contrast, mice that received the primary infection of SARS-CoV-2 followed by RSV demonstrated worsening SARS-CoV-2-induced disease with no alteration of SARS-CoV-2 replication. Interestingly, these mice had simultaneous protection from RSV-induced disease and reductions in RSV replication. Collectively, these findings suggest that exposure to RSV before SARS-CoV-2 may provide protection from SARS-CoV-2 induced disease, while exposure to RSV after SARS-CoV-2 could potentially worsen SARS-CoV-2 pathology.

**Fig. 5:**
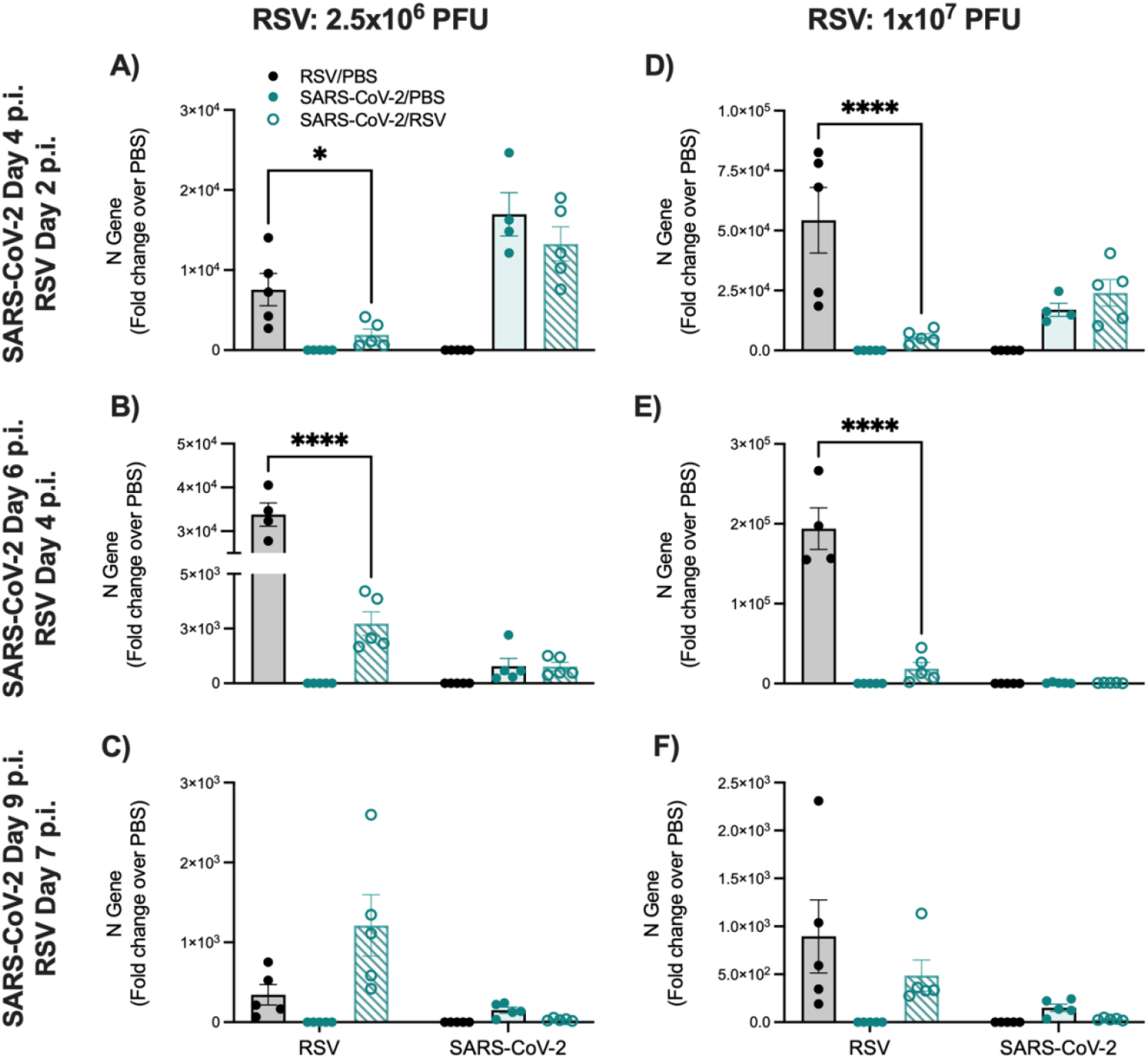
Assessment of gene expression by RT-qPCR in the lung of BALB/c mice infected with SARS-CoV-2 followed by RSV. Mice were infected as shown in Figure 3A. Lung tissue was collected at (A,D) day 4 post-SARS-CoV-2 infection/day 2 post-RSV infection, (B,E) day 6 post-SARS-CoV-2 infection/day 4 post-RSV infection, and (C,F) day 9 post-SARS-CoV-2 infection/day 7 post-RSV infection to assess RSV N and SARS-CoV-2 N gene expression by RT-qPCR. Data are expressed as mean ± SEM. Significant results as compared to the respective controls are marked with asterisks (* p ≤ 0.05, **** p ≤ 0.0001).

These changes in clinical disease and viral replication are likely influenced by viral-viral interactions such as direct viral interference or inhibition of the secondary virus by the host response to the primary infection. In our models of simultaneous RSV/SARS-CoV-2 co-infection and secondary SARS-CoV-2 infection, replication of SARS-CoV-2 in the lung tissue is significantly reduced. This could be a consequence of RSV-induced killing of epithelial cell types key to the progression of SARS-CoV-2 from the upper to lower airways [17-20]. Additionally, priming of inflammatory cytokines and type-I interferons (IFN-I) have shown to have antiviral effects during SARS-CoV-2 infections in vitro [21]. Though RSV is thought to be a poor inducer of IFN-I when compared to other respiratory viruses, IFN-I is still induced by RSV infection along with a strong TNF-α and IL-6 response [22]. Therefore, inflammation induced at early timepoints and downstream activation of interferon stimulating genes (ISGs) could contribute to the limiting of SARS-CoV-2 replication in both our simultaneous and secondary co-infection models. This is supported by a recent study which shows the suppression of SARS-CoV-2 in a model of simultaneous RSV/SARS-CoV-2 co-infection in human bronchial epithelial cells (HBECs) to be mediated by ISG15 and IRF-3 signaling [23].

Interestingly, primary infection with SARS-CoV-2 followed by RSV resulted in reduced RSV replication in the lung. Similar mechanisms of epithelial damage could potentially explain this reduction, limiting RSV replication to the upper airways [18]. Since IFN-I expression has been previously shown to not directly alter RSV replication, it is more likely that the immune cell repertoire elicited at day 2 p.i. to SARS-CoV-2 infection may be ideal for limiting RSV replication [24]. Additionally, these SARS-CoV-2/RSV infected mice had worsening SARS-CoV-2 induced clinical disease. This could be explained by an increase in cytokine activity or ISG induction by RSV that could further impact the inflammatory response, resulting in worsening of SARS-CoV-2 pathology [25]. Our findings of inhibition of the secondarily infected virus are consistent with what has been shown for SARS-CoV-2/Influenza co-infections in mice [15].

Interestingly, our model of RSV/SARS-CoV-2 differs from SARS-CoV-2/Influenza co-infection in that the latter leads to prolonged replication of either primarily infected virus. Our data would suggest that the primary infecting virus is either unaffected or cleared from the lung more efficiently (Figure 4). Having established the baseline dynamics of clinical disease and viral replication through our co-infection model, future characterization of the immune response and viral spread during RSV/SARS-CoV-2 co-infection should be investigated.

At the beginning of the COVID-19 pandemic, cases of SARS-CoV-2 among children were surprisingly rare [2]. It wasn’t until the historic low in circulation of other respiratory viral infections during the 2020/2021 winter season, and the emergence of the Delta variant in mid-2021 that we began to see more consistent SARS-CoV-2 infections in the pediatric population [2-4].Interestingly, the 2019/2020 RSV season was described as having a high number of RSV infections and lasting longer than usual in some countries [26-28]. Based on the data presented in this study, it is plausible that RSV infections among children during the 2019/2020 season could have contributed to this resistance to infection during the initial 2019/2020 SARS-CoV-2 wave. With the anticipated release of the RSV vaccine in the coming months, it is also important to consider how this may affect the RSV/SARS-CoV-2 interaction. Pfizer has reported that their RSV vaccine is 81.8% effective at reducing the occurrence of severe RSV infections in infants [29]. Our data would suggest that having even a mild RSV infection would potentially protect from SARS-CoV-2 induced disease.

Taken together, this study has established a working animal model for and provides important insights into RSV/SARS-CoV-2 co-infections. Our findings suggest that exposure to RSV before SARS-CoV-2 may provide protection from SARS-CoV-2 induced disease, while exposure to RSV after SARS-CoV-2 could potentially worsen SARS-CoV-2 pathology. Moreover, this study highlights the potential impact of RSV infections on the susceptibility and resistance to SARS-CoV-2, especially in the pediatric population. Further research is needed to better understand the underlying mechanisms driving these changes in clinical disease and viral replication. Given these baseline characteristics, our model could be used to strengthen treatment strategies for co-infected patients and to further our knowledge of the unique interplay between RSV and SARS-CoV-2.

## Materials and Methods

### Ethics statement

All care and procedures involving mice in this study were completed in accordance with the recommendations in the *Guide for the Care and Use of Laboratory Animals* of the National Institutes of Health and the UTMB institutional guidelines for animal care. The Institutional Animal Care and Use Committee (IACUC) of UTMB approved these animal studies under protocol 2102014.

### Virus preparations

The mouse-adapted SARS-CoV-2 strain, CMA3p20, was a gift from Dr. Vineet D. Menachery. Information pertaining to the development of CMA3p20 can be found here [30].Propagation of CMA3p20 was done as described previously [31]. All mention of SARS-CoV-2 in this study pertains to the use of CMA3p20. All virus preparations for SARS-CoV-2 were performed by trained personnel in a biosafety level 3 (BSL-3) facility.

### Animal infections

Female, 8 to 10-week-old BALB/c mice were purchased from Envigo (Indianapolis, IN, USA) and maintained in Sealsafe HEPA-filtered air in/out units. For infection, mice were anesthetized with isoflurane and infected intranasally (IN) with SARS-CoV-2 and/or RSV diluted in 50uL of PBS. For SARS-CoV-2 infections, mice were IN inoculated with a dose of 1×10^6^ TCID_50_/mL. For RSV infections, mice were IN inoculated with 2.5×10^6^ PFU or 1×10^7^ PFU of RSV Long Strain. For simultaneous co-infection experiments, SARS-CoV-2 was combined with RSV in PBS prior to inoculation. For co-infection 48 h apart, mice were IN inoculated with either SARS-CoV-2 or RSV day 0. On day 2, mice were anesthetized and then IN inoculated with the corresponding virus. All animals were monitored for weight loss and illness was scored as described [32]. At days 2, 4, 6, 7, and 9 p.i., lung tissue was collected for assessment of viral replication by RT-qPCR. Due to the lack of weight loss in the 2.5×10^6^ PFU RSV mice, lung tissue was collected at day 4 p.i. and active infection was confirmed by plaque assay as described (data not shown; [33]). All animal experiments involving infectious virus were performed in UTMB’s animal biosafety level 3 (ABSL-3) facility by trained personnel with routine medical monitoring of staff.

### Assessment of Viral N Gene by RT-Qpcr

The right lung was collected at the timepoints indicated above. Tissue was homogenized in TRizol and 500μL of tissue lysate was subjected to phase separation using chloroform. The top aqueous layer was then further processed using the Qiagen RNeasy Mini kit in accordance with the manufacturer’s instructions. Isolated RNA was directly subjected to one-step RT-qPCR analysis using TaqMan Fast Virus 1-Step Master Mix (Thermo Fisher Scientific, MA, USA) and Bio-Rad CFX instrumentation (Bio-Rad, CA, USA). The following custom TaqMan gene expression assay IDs were used to assess the expression of RSV N and SARS-CoV-2 N genes: ARU66XH and APNKYWD (Applied Biosystems, CA, USA). A no template control was included in each run. One-step RT-qPCR reactions were run as follows: 50C for 5 min, 95C for 20s, followed by 40 cycles of 95C for 15s, then 60C for 60s. Cycle threshold (C_T_) values were analyzed in Microsoft Excel by the comparative C_T_ (ΔΔC_T_) method according to the manufacturer’s instructions (Applied Biosystems). The amount of target (2^−ΔΔCT^) was obtained by normalization to the endogenous reference (18S) sample. Fold change in gene expression was calculated in comparison to the PBS control mice.

### Statistics

Statistical analyses were performed using an ordinary two-way ANOVA followed by Tukey’s multiple comparison test, a mixed-effects model followed by Geisser-Greenhouse correction, or an unpaired student’s t-test (GraphPad Prism 9.5.1; GraphPad Software, Inc., San Diego, CA, USA). Results are expressed as mean ± SEM for each experimental group and p ≤ 0.05 value was selected to indicate significance.

## Acknowledgements

We would like to thank Slobodan Paessler for his assistance with the BSL3 facilities and Michelle N. Vu for her helpful discussions. The graphical figures were created using BioRender.com.

## Contributions

DRM conceptualized this study, obtained funding for this study, conducted the animal experiments, and wrote the manuscript. YQ conducted the animal experiments and edited the manuscript drafts. KST and AHM conducted the RT-qPCR experiments and edited the manuscript draft. RP created the graphical abstract. VDM provided the mouse-adapted SARS-CoV-2 virus. AC and RPG obtained funding for this study and provided experimental support.

## Funding

This research was funded by UTMB Institution for Human Infections and Immunity (IHII) Data Acquisition Grants as well as the NIH grant AI062885.

## Notes

### Competing Interest Statement

The authors have declared no competing interest.

